# A Screen Reveals Bexarotene, Sorafenib and Certain of its Analogues as Common and Effective Inhibitors of Growth of Bacteria that Grow on LB but not on McConkey Agar

**DOI:** 10.1101/2023.04.01.535200

**Authors:** J. Scheerer

## Abstract

A screen of 61 compounds identified bexarotene and sorafenib as inhibitors of bacterial isolates that grew on LB agar but not on McConkey agar. The effect of bexarotene was durable, whereas sorafenib simply retarded growth. Two sorafenib analogues, previously not described, dubbed JSR-A40 and JSR-N42, displayed durable effects. Upon extending this to a larger group of bacteria from different sources it was found that both bexarotene and JSR-N42 were active in concentrations from between 1.25-10 µM in 42 of bacteria growing in LB but not McConkey medium, whereas bacteria growing on McConkey agar were all found to be insensitive at that concentration. Both bexarotene and N42 displayed rather broad characteristics similar to clindamycin, however non-overlapping. Sensitivity to clindamycin, if observed, was generally higher. It was found that the urea type compound N42 at 5 µM was broader active than clindamycin at 5 µM in this set of isolates.

## Introduction

The screening of chemical compound libraries is a mainstay for the identification of drugs and the validation of putative targets. Examples of recent developments in antibiotics research include drug repurposing [1, 2], computational methods [3], and tapping previously unaddressed sources [4]. In yeast, genetic interaction profiling as well as the search for interacting drug pairs are means to analyse complex networks [5, 6] for exploitation in drug discovery [7] and combining approaches did help to substantiate concepts such as synthetic lethal interactions [cf. 8,9] on which a highly successful anticancer drug emerged [10]. However, given the tremenduous complexities associated with such research any additional information should prove helpful. Therefore, it was undertaken to screen a 61 compound collection for possible growth effects in yeast. During these experiments it was found that not yeast but certain contaminations displayed sensitivity to two compounds in that set, bexarotene and sorafenib. From extending these findings to a broader set of isolates it is reported here that both bexarotene and certain urea compounds based on a Sorafenib scaffold offer a broad activity in bacteria which based on their failure to grow on McConkey agar would be considered gram-positive. The possible clinical relevance is discussed.

## Materials and Methods

All bacteria were considered non-fastidious and collected and diluted in Lennox medium as needed and grown on Lennox agar supplemented with 20% of its weight of YPD, or on McConkey agar, as recommended. For assay purposes a number of up to 30-100 colony forming units per 15 µL applied was found to yield best results. Drugs were dissolved in DMSO and were stored at 4°C. For drug containing plates an appropriate amount of drug dissolved in DMSO was added to the molten agar at temperatures of 60-70°C and stirred on a magnetic stirrer until evenly distributed. Plates were allowed to dry at ambient temperature and moisture for 5-7 days to enable both drying for rapid absorption of bacterial suspensions and equilibration of drug and DMSO. Weight losses were typically 200mg/day and these resulted, as was found and expected, from the top third volume of a 20 mL plate. All media were from Roth, Karlsruhe, Germany.

Error estimates were derived from the Einstein-Smoluchowski equation with diffusion coefficients for small molecules in agarose gels. This suggested that the average travel distance is large compared to the thickness of the layer and therefore, DMSO and drug should evenly distribute within 3-5 days. Likewise, growth of bacteria at room temperature should gradually effect equal distribution of DMSO and drug into the volume applied.

The assessment of growth on agar plates was by visual inspection of agar plates either as photographed or in translucent light after 3-10 days time at ambient temperature to allow for colonies to form on control plates devoid of drug. The concentrations given here as “inhibitory concentration” are those were no visible colonies were detectable even after 4 days or longer. Growth in liquid culture was assessed by serially diluting four times one hundred-fold and comparing densities at the respective dilutions versus control and versus time zero. Zero growth indicated full growth inhibition in the presence of drug with no colonies visible. Zero growth also hinted at possible bactericidal effects.

Compounds investigated are listed in appendix A. They were either obtained from pharmaceutical material (PH) and worked up and crystallized as appropriate or were purchased from Chemietek, Indianapolis (CT). Bexarotene was both from CT and from ABCR, Karlsruhe, Germany. Others were obtained by custom syntheses (CS) according to published procedures available to those skilled in the art. Identities and purities were verified by HPLC and NMR spectroscopy. The stereochemistry of spirooxindole MI-63X was assessed by NMR HSQC and HMBC coupling experiments. HPLC assessments were done by former Infraserv Knapsack now Synlab. NMR measurements were done by Dr. E. Haupt of Department of Chemistry, University of Hamburg, Hamburg, Germany. Mass spectroscopy and preparative HPLC, as needed, was done by Fischer Analytics, Bingen, Germany.

Synthesis of sorafenib analogues: Briefly, 10 mmol of 4-chloro-3-trifluorophenylisocyanate was dissolved in 15 mL of toluene. One equivalent of amine suspended or dissolved in 15 mL of toluene was added and stirred at ambient temperature for 16 hours. The resulting crystals were filtered dry and washed with cold toluene. The corresponding amine-hydrochlorides were suspended in 10 mL dichloromethane and added to the isocyanate solution. Then, 1 equivalent of DIPEA dissolved in 5 mL dichloromethane was added under stirring. After stirring for 16 hours at room temperature the volume was concentrated to remove the dichlormethane and washed with water. Typically, crystals formed within hours and were isolated and suctioned dry. The analogues were used without further purification. Compound A40 used 2-adamantanamine as the amine (Aldrich; CAS10523-68-9) and N42 used 4-amino-3-nitrophenol (ABCR, CAS 121-88-0). Another compound, P43, used pyrrolidine (Fluka) for coupling, T44 used 3,4,5-trimethoxyaniline (ABCR, CAS 24313-88-0) and T45 DL-Trp-ethylesterhydrochloride (Aldrich, CAS 6519-67-1). The urethane E41 used ethanol as the alcohol and was prepared by reacting the isocyanate with ethanol in dichloromethane.

## Results

A screen aimed at investigating the response of ordinary bakers yeast to a number of 61 chemicals of pharmaceutical interest was performed. A third of these were kinase inhibitors (see appendix A). It was noted that not yeast, but 6 nonidentified contaminants, likely of bacterial origin, reproducibly responded to two drugs from the library. These, bexarotene and sorafenib, in 4 of these 6 contaminants resulted in inhibition with no growth visible in the single digit micromolar range and in dose-dependent fashion. None of the other compounds did reach a 10 µM limit.

To investigate whether this could be reproduced elsewhere, a total of 11 isolates from the human body surface, 4 off white ones from the anal region, 4 off white ones from scrotal region, 3 yellow ones from the neck area were cultivated alongside with 5 random picks that were obtained from older plates. Attempts to obtain isolates from armpit, mouth and navel on LB agar failed. This group of isolates was dubbed "ELSA“, and they were extensively used to further investigate behavior of bexarotene and sorafenib on their growth. In the agar dilution test used it was found that bexarotene was able to fully suppress growth to zero visibility at concentrations of 10 µM in these isolates and this suppression was maintained for at least 14 days. On contrast, sorafenib appeared slightly more potent on a short time scale, but inhibition appeared short lived and typically, after about a week one would not have considered sorafenib to be effective. In the assay used it was easy, from measurement of colony diameters, to calculate the extent of inhibition at some time point, as drug induced halving or quartering of the radii of colonies amounts to about 88% and 98%, resp., growth inhibition at that time point. While this was true and found for sorafenib on a shorter time span, colony growth carried on, however, until no visible difference versus control was discernible.

A literature search then showed that sorafenib had been earlier identified as being active against staphylococci, displaying similar characteristics as observed here and highly effective analogues had been synthesized [1, 10]. Therefore, from the precursor isocyanate of sorafenib, a number of simple analogues was made from off the shelf amines, using standard methods for urea syntheses (see Materials and Methods). Of 6 of these analogues two, a compound with a very bulky substituent dubbed JSR-A40 and a nitrogroup containing compound dubbed JSR-N42 (see Material and Methods) much to surprise were found to be more effective than sorafenib at same concentrations. In particular, efficacy was maintained and at 10 µM no (N42) or almost no growth (A40) was detected in ELSA isolates on agar plates, even after a week, quite similar to bexarotene. Compound A40 at 10 µM displayed very little growth in a few of the ELSA isolates after days of incubation, however, N42 shut off growth for all of them at 10 µM. As the bulky substituent of A40 is also present in a similar urea-type drug, AU1235, this was also tested, and it was found that this compound at 20 µM displays activity identical to negative control (not shown). Thus, substituting the sorafenib halogen containing group against a 2,3,4-trifluorophenyl containing core rendered an active compound inactive and any structure activity related research should probably focus on the opposite side of the urea moiety. In line with findings reported earlier [10], 3,4,5-trimethoxyaniline as a coupling partner was ineffective and so was D,L-tryptophanethylester in all of the ELSA isolates. 2 other coupling partners, pyrrolidine and ethanol, resp., were designed to be ineffective controls and proved to be so.

From liquid culture experiments comparing bexarotene, A40, N42 and sorafenib it was estimated that bexarotene and N42 reproducibly displayed highest activity (not shown). When select members of the ELSA group of bacteria were grown in the presence of 20, 6.6 or 2.2 µM of bexarotene or N42 it was found that at the highest concentrations no growth at all was observed or was attenuated by at least 3-4 and up to 6 orders of magnitude versus negative control (appendix B). When isolate E1 was incubated in the presence of 5 or 10 µM of N42 it was found that 10 but not 5 µM of N42 reduced the number of visible colonies versus starting numbers (Table 1a). This was not further investigated. With 5 to 10 µM being the most likely range for eliciting a major dent in the ability to grow a concentration 7.5 µM was included in a kinetics experiment on isolate S3. It was found that after 54 h of incubation isolate S3 in the presence of bexarotene at 10, and N42 at 7.5 µM suppressed growth to an extent that no development was recorded (Table 1b). The relative difference between bexarotene and N42 was established by serially diluting samples in that experiment after 78 h of incubation and comparing their numbers. It was found that Bexarotene displayed a rather sharp decline in density between 7.5 and 10 µM and albeit N42 at 5 µM displayed a lawn when not diluted, the difference in efficacy is close to a factor of two as suggested by agar dilution assays (Table 1c).

**Table 1.** Kinetics of Bexarotene and N42 at Various Concentrations on Isolates E1 and S3.

**a) N42 kills isolate E1 at 10 but not at 5 μM.**
| Time [h] | Number of colonies per spot (15 μL) | |
| --- | --- | --- |
| | N42 5 μM | N42 10 μM |
| 0 | 17/18/16/10 | 13/16/18/10 |
| 9 | 9/16/24/18 | 12/11/13/9 |
| 24 | >200/100/90/80 | 0/5/2/0 |
| 48 | Lawn/lawn/lawn/lawn | 0/0/0/0 |

**b) Effect of Various Concentrations of Bexarotene and N42 on Isolate S3.**
| Time [h] | Colony numbers per 15 μL in the presence of drug | | | | | |
| --- | --- | --- | --- | --- | --- | --- |
| | Bexarotene [μM] | | | N42 [μM] | | |
|  | 10 | 7.5 | 5 | 10 | 7.5 | 5 |
| 0 | 41 | 45 | 36 | 30 | 34 | 31 |
| 3 | 40 | 31 | 36 | 27 | 28 | 36 |
| 9 | 42 | 38 | 27 | 22 | 15 | 29 |
| 18 | 30 | 57 | 69 | 24 | 32 | 33 |
| 30 | 73 | Lawn | Lawn | 32 | 25 | $\geq 350$ |
| 42 | 34 | Lawn | Lawn | 20 | 77 | Lawn |
| 54 | 70 | Lawn | Lawn | 26 | 132 | Lawn |

| Dilution<br>Factor<br>(1:33) | Colony numbers per 15 µL after 78 h |  |  |  |  |  |
| --- | --- | --- | --- | --- | --- | --- |
|  | Bexarotene [µM] |  |  | N42 [µM] |  |  |
|  | 10 | 7.5 | 5 | 10 | 7.5 | 5 |
| 0-fold | >200 | Lawn | Lawn | >200 | >200 | Lawn |
| 1-fold | 12 | Lawn | Lawn | 10 | 29 | Lawn |
| 2-fold | 0 | Lawn | Lawn | 0 | 0 | ~150-200 |
| 3-fold | 0 | Lawn | Lawn | 0 | 0 | 22 |
| 4-fold | 0 | ~150-200 | Lawn | 0 | 0 | 10 |

In a next step it was checked whether the ELSA bacteria would grow on McConkey agar. This was not the case and they were thus considered gram-positive. It was concluded that bexarotene and N42 effectively inhibited growth of 16 gram-positive bacteria at concentrations between 5-10 µM. From a repetition experiment that included clindamycin at 10 µM, it was concluded, by comparison, that the ELSA group likely could be classified as a group of just 5 bacterial species. Mutual exclusivities were found, i.e. bexarotene inhibited growth where clindaymcin did not and vice versa in the S and A group as well as L4 and L5 (compare Table 3, Plate D).

To further investigate whether these findings could be extended to bacteria isolated from other sources, samples were collected from the bathroom sink, from a backyard water pitch, from suburb garden soil, from a creek mud bank and from a cattle conditioned meadow as well as 6 on McConkey growing isolates from human faeces (Table 2). These all were checked for growth on McConkey agar and tentatively classified as gram-positive if they did not. It was found that gram-negative McConkey growing bacteria were not susceptible to neither of the compounds sorafenib, A40, N42, bexarotene nor clindamycin at 10 µM. In a set of McConkey nongrowers, however, the very same pattern as already expected, was observed again. Bexarotene was effective at 10 µM and inhibition was maintained, as it was in the presence of A40 and N42, however, less so in the presence of sorafenib, whereas the analogues made from trimethoxyaniline and Trp-Ethylester were as ineffective as was AU1235 (not shown).

**Table 2.** Sensitivities to Sorafenib, A40, N24, Bexarotene and Clindamycin of Bacteria from Various Sources Grown on McConkey Agar

| Source | Number of colonies picked | Primary selection medium | Number growing on McConkey agar | Growing on McConkey and growing in the presence of either of 5 drugs at 10 μM | Estimate of number of different species in sample picked |
| --- | --- | --- | --- | --- | --- |
| Faeces | 6 | McConkey | 6 | 6 | 2 |
| Sink | 8 | Lennox | 8 | 8 | 5-7 |
| Soil sample 1 | 8 | Lennox | 1 | 1 | n.a. (>1) |
| Soil sample 2 | 6 | Lennox | 4 | 4 | n.a. |
| Soil sample 3 | 8 | Lennox | 8 | 8 | 3 |
| Creek bed | 8 | Lennox | 3 | 3 | n.a. |
| Cattle conditioned meadow | 9 | Lennox | 1 | 1 | 6-7 |

31 out of the 53 colonies that were initially picked from the sources described above grew on McConkey agar and 22 did not grow on McConkey agar.
All bacteria that grew on McConkey also grew in the presence of all of either of the 5 drugs Sorafenib, Bexarotene, A40, N42 and Clindamycin at a
concentration of 10 μM with no growth rate difference as judged by comparison with a blank control. N.a. not assessed. The estimate of species is
based on shape and color.

These experiments were then taken further by including all those bacteria that remained as classified gram-positive. 46 bacteria, out of which 4 were included that grew on McConkey agar, were subjected to varying concentrations of bexarotene (2.5; 5 and 10 µM), clindamycin (1.25; 2.5; 5 and 10 µM) and N42 (1.25; 2.5; and 5 µM) in the presence of 2.5% DMSO or 5% DMSO. Bacillus cereus was omitted due to its peculiar growth characteristics. It had separately been tested sensitive to A40 and N42, though. One other, which released a deep brown color was omitted as well due to adverse cultivation characteristics. Still another one, believed to be Rotundula, a yeast, proved to be insensitive.

The results are compiled in Table 3 for the 2.5% DMSO experiment and they are not different from the 5% DMSO experiment which is not shown. It can be seen that 4 McConkey growers in accordance with the previous findings did not respond to any of the drugs at any of the concentrations used. One germ that was equally non responsive to all drugs was, however, a McConkey nongrower. Generally, as was expected from the experiments conducted so far, N42 was broadly active even at 5 µM and thus slightly more active than bexarotene, which needed 10 µM to do so. Clindamycin however, was the most effective compound in all those cases were it worked (Table 3).

**Table 3.**
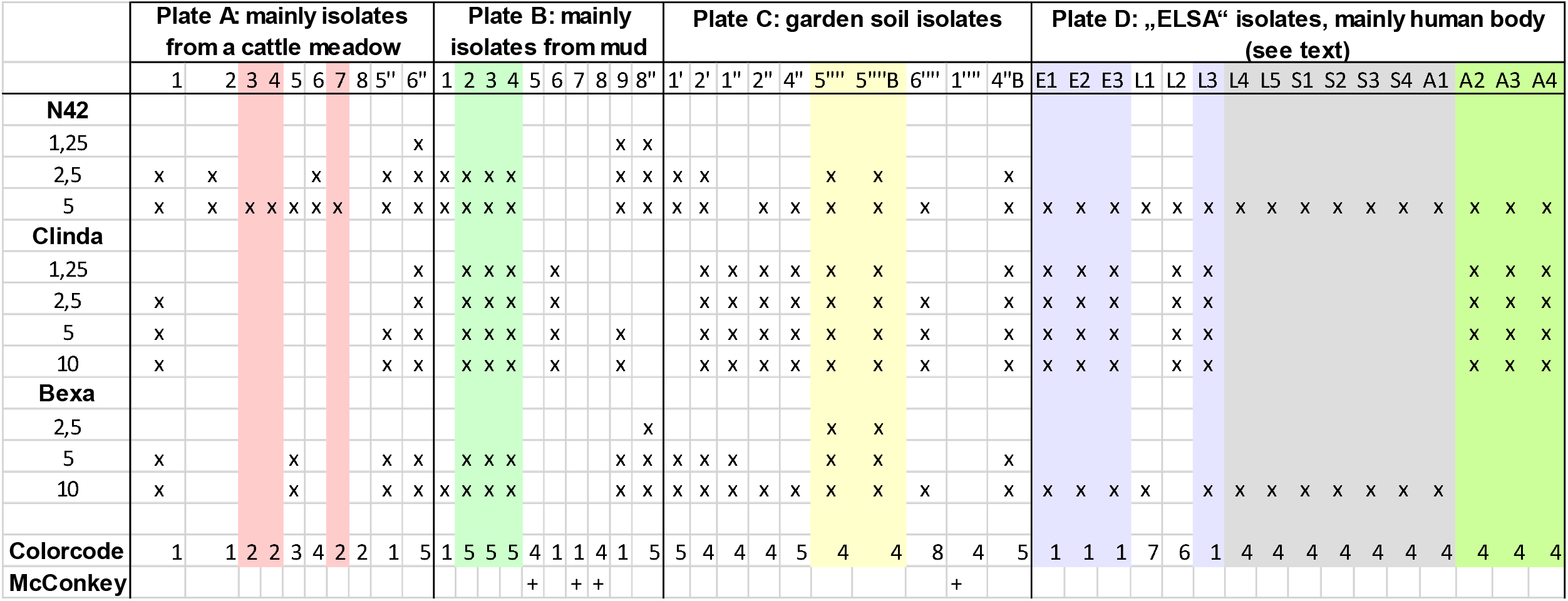
Agar Dilution Assay Results of N42, Clindamycin and Bexarotene in 46 Bacterial Isolates. Results of various concentrations of JSR-N42, Clindamycin and Bexarotene. Concentrations given in μM DMSO concentration is 2,5%. A parallel experiment containing 5% DMSO was conducted and displayed identical results. Colorcode: 1=yellow; 2=orange; 3=light orange, apricot; 4=off -white; 5= white; 6=plain/bright white; 7=greenish-yellow; 8=whiteish yellow. The colors of the shaded areas indicate apparent identities as judg ed from these criteria. Drug concentrations are in mikroM. The marks (“X”) indicate no visible growth of colonies by time enough to have clear evidence of v isible colonies on control plates (3-5 days at room temperature). No visibility means no colonies visible neither on a photograph nor by looking through the plat e. Colony 8 on plate A was verified as a McConkey-Nongrower, as it is the only germ in this set that behaves like a McConkey Grower. “+” defines bacteria that grew on McConkey agar.

From varying concentrations of DMSO it was clear that increasing DMSO increasingly induced growth defects. It was therefore, attempted to lower DMSO concentrations as far as possible. It was found that bexarotene, as a highy hydrophobic compound could be administered without loss of activity to molten agar at 60-70°C to achieve a final concentration of 2.5% DMSO, as no difference was detectable versus 5%, and all expected patterns remained, irrespective of DMSO concentration. It was found, however, that 2 neighboring colonies grew to better visibility on the 5% DMSO than on the 2.5% DMSO plate in the presence of 5 µM bexarotene, whereas all other colonies behaved as expected. The reason for this remains unelucidated and the experiment was not repeated as the overall dose-response relation was not affected. It should be remarked, however, that a similar counterintuitive behaviour had been observed in 2 separate experiments, one including bexarotene (see Appendix B, isolate A1) and another including A40, a similarly hydrophobic compound (not shown). N42, in contrast to bexarotene is soluble in water to at least 100 µM, yet distributes very well into octanol, and thus does not require DMSO to be dissolved. It was also noted that some, but not all of the group of bacteria investigated were rather sensitive to even 5% DMSO. This did not however, impact on the outcome of clindamycin, bexarotene or N42 or on the overall pattern. It was not investigated whether there were any synergies between DMSO and any of these drugs.

The 42 McConkey-nongrowing bacteria were finally classified based on their responsiveness to varying concentrations of bexarotene, N42, clindamycin, their appearance and their color. After curation it was estimated they probably consisted of 26 independent McConkey sensitive isolates (Table 3). Assuming this assessment is correct, then 23, 16 and 12 different species, resp., out of a total of 26 were sensitive towards 5 µM of JSR-N42, clindamycin and bexarotene resp., to an extent they did not form any visible colonies versus negative control after 7 days of incubation at room temperature.

## Discussion

These experiments revealed that the efficacy of certain sorafenib analogues, bexarotene and clindamycin was restricted to bacteria that do not grow on McConkey agar and would therefore be considered gram positive. While clindamycin did not result in zero growth of all of the selected group even at 10 µM, it was possibly to achieve this with either bexarotene or JSR-N42 even at 5 µM, except in one single case. However, if clindamycin achieved zero growth it was at concentrations down to 1.25 µM.

An agar dilution assay was used to determine sensitivities. A single day at ambient temperature and moisture leads to loss of approx. 200 mg of weight from a plate and this is restricted mainly to the surface. Thus, this might induce some concentration gradient in drug or DMSO over a few days (see Materials and Methods). When 15 µL of bacterial suspension then was added this might have led to some local dilution again. Therefore, the two counterbalancing effects may have induced some error in the actual concentrations. As the effect of dilution from adding the bacterial suspension should be comparatively large in comparison to diffusion effects that result from drying the plate it may well be that the efficacies are somewhat higher than indicated. It was noted also, though, that the density of administered suspension has an effect on the reading, as described elsewhere [12]. Higher densities bias towards lower efficacy. This may be due to either local target saturation or some sort of quorum sensing or both. It was found safe to obtain a streak from a plate, the size of the tip of a loophole, dilute this into 1mL medium and dilute this twice by 1:100 to achieve highly reproducible outcomes, with not even duplicates being actually necessary.

The compound JSR-N42 is a very simple drug that contains a free phenolic hydroxyl that should be amenable to being modified for affinity chromatography. As this type of compound is actually a precursor for sorafenib or regorafenib type syntheses, it is clear that the additional groups that exist in these latter drugs are not necessary for efficacy in bacteria and therefore N42 probably is safe for modification on the hydroxyl.

The clinical relevance of these findings are probably limited, as there are many excellent antibiotics available and only MRSA represent a major threat in the hospital. Bexarotene is licensed in man for cutaneous T-cell lymphoma and from the pharmacokinetics outlined in its SmPC it seems plausible that concentrations of 10 µM can be reached and maintained in the human body. JSR-N42, although equally or more effective than bexarotene, is not a licensed drug and despite apparent attractiveness as regards its solubility, the presence of an unmodified hydroxyl would render it susceptible to rapid glucuronidation and excretion. In addition, full scale clinical development would be necessary for use in man.

A weakness of the present study is that the species have not be characterized and no information on efficacy in biofilms has been generated. However, the identification of these two compounds may help with research into this.

Both compounds however, should be suitable to adress targets in gram-positive bacteria that may help to further understand the intricate networks inside a cell. There exist clues as to what these targets may be in the case of N42 (references cited in [10]). Bexarotene, on the other hand, has not, to the best of our knowledge, been described previously as effective in bacteria and it should be interesting to figure out what its target turns out to be.

In conclusion, results on sorafenib analogue compounds have been corroborated [1, 10]. Two novel analogues, JSR-N42 and JSR-A40 in addition to those already described have been synthesized. The retinoic acid agonist bexarotene has been identified as a new inhibitor. The results show these compounds are active in a broader range of gram-positive bacteria.

## Appendix A

### List of Compounds

**Source description: CS, Custom synthesis as described in materials and methods; CT, Chemietek, Indianapolis, USA; PH, pharmaceutical source**

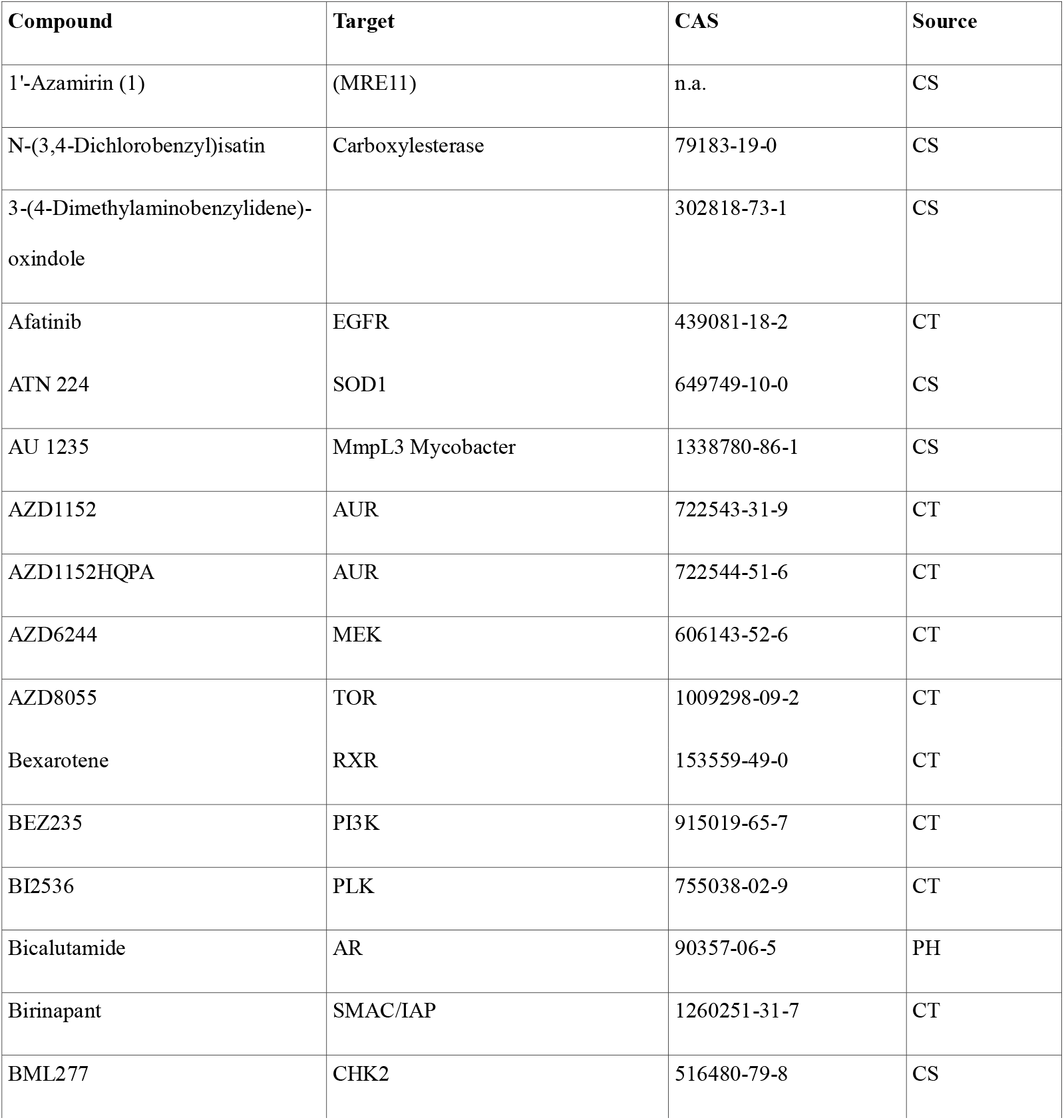
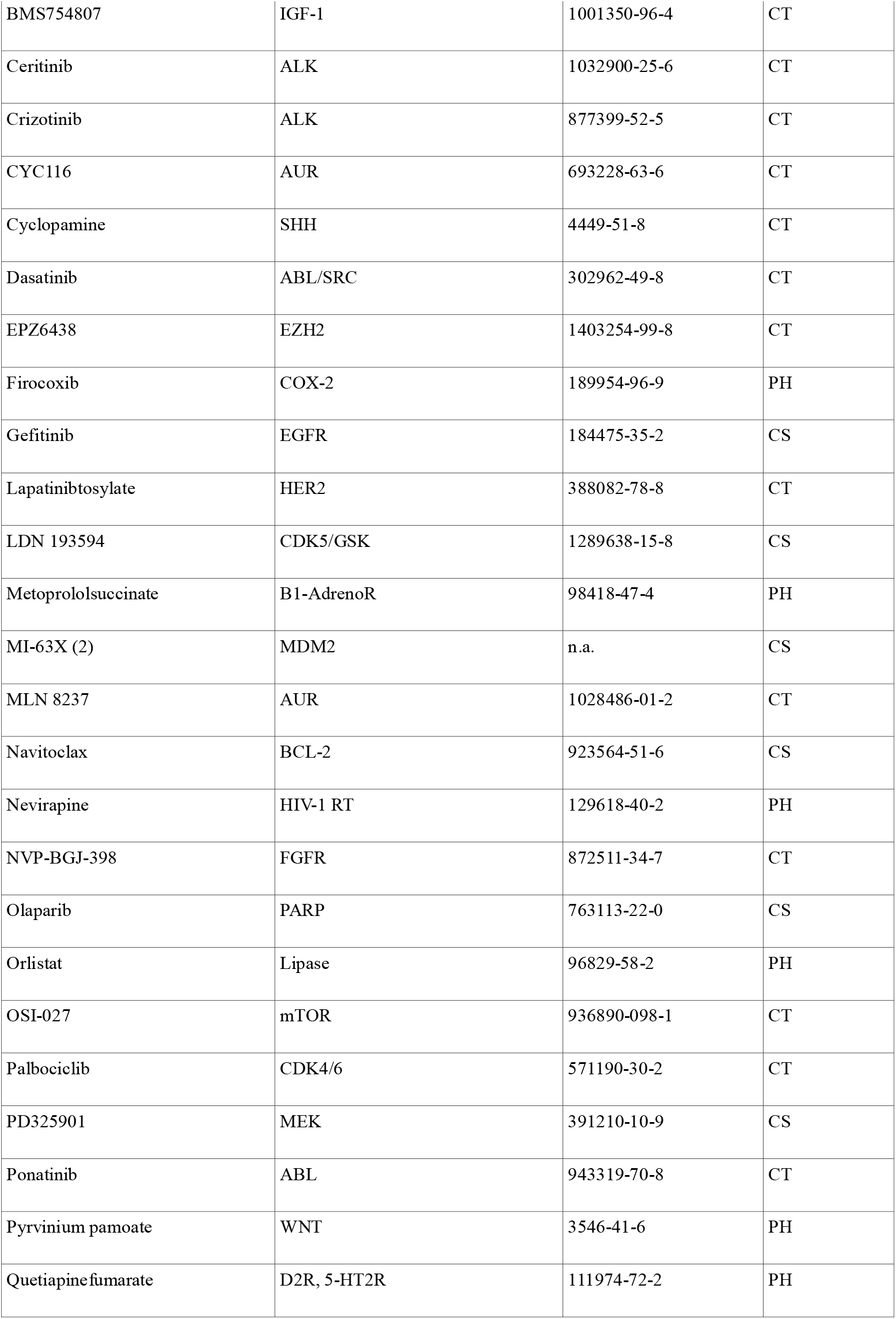
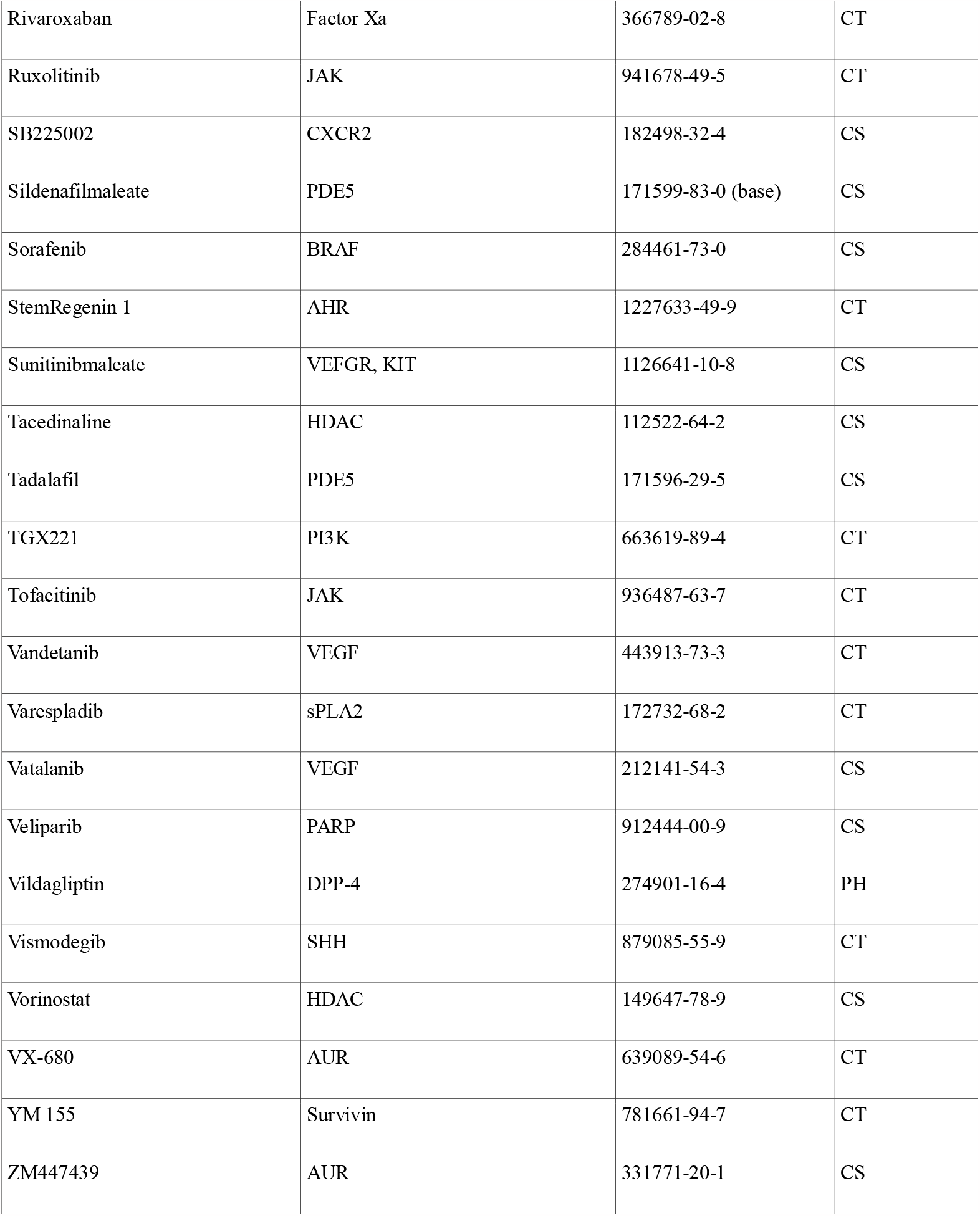

1. 1’-Azamirin is a derivative of Mirin, which contains Nitrogen in the 6-membered ring and has been formed using 4-Pyridylaldehyde instead of 4-Hydroxybenzaldehyde in the Pseudo-Thiohydantoin condensation reaction. Dupre A, Boyer-Chatenet L, Sattler RM et al., A forward chemical genetic screen reveals an inhibitor of the Mre11-Rad50-Nbs1 complex. Nature Chem Biology. 2008 4(2), 119-125. Doi:10.1038/nchembio.63. Garner, KM, Pletnev AA and Eastman, A. Corrected structure of mirin, a small molecule inhibitor of the Mre11-Rad50-Nbs1-complex. Nature Chem. Biology 2009, 5(3). Doi: 10.138/nchembio0309-129.
2. Spirooxindole MI-63X is a hybrid of spiroxindoles MI-63 and MI-219, where the halogens in the ring system are from MI-219 and the morpholinoethylamino-sidechain is from MI-63. Ding K, Lu Y, Nikolovska-Coleska Z, et al., Structure-based Design of Spiro-Oxindoles as Potent, specific Small-Molecule Inhibitors of the MDM-p53 Interaction, J.Med.Chem. 2006, 49, 3432-3435. Ding K, Wang G, Deschamps J, et al. Synthesis of Spirooxindoles via Asymmetric 1.3-dipolar Cycloaddition: Tetrahedron Letters 2005, 36, 5949-5951. Beloglazkina A, Zyk N, Majouga A etal. Recent small-Molecule Inhibitors of the p53-MDM2 Protein-Protein Interaction. Molecules 2020, 25, 1211; Doi: 10.3390/molecules

## Appendix B

**Effects of Bexarotene and N42 toward Select Members of the ELSA Group of Bacteria in Liquid Culture**.

| Isolate | Compound | Density versus Negative Control at Concentration given (μM) |  |  |
| --- | --- | --- | --- | --- |
|  |  | 20 | 6.6 | 2.2 |
| E1 | Bexa | 0 | 1 | 1 |
|  | N42 - #1 | 0 | 1:10,000 | 1 |
|  | N42 - #2 | 0 | 0 | 1 |
|  | N42 - #3 | 0 | 0 | 0 |
|  | N42 - #4 | 1:10,000 | 1:100 | 1 |
| A1 | Bexa - #1 (8%) | 1 | 1 | 1 |
|  | Bexa - #2 (2%) | 1 | 1 | 1 |
|  | Bexa - #3 (5%) | 0 | 1 | 1 |
|  | Bexa - #4 (5%) | 1:1000 | 1 | 1 |
|  | N42 - #1 (8%) | 0 | 0 | 1 |
|  | N42 - #2 (2%) | 0 | 0 | 1 |
| S4 | Bexa | 1:10,000 | 1 | 1 |
|  | N42 | 1:10,000 | 1:1,000-10,000 | 1 |
| L1 | N42 | 0 | 1:10,000 | 1 |
| L2 | N42 | <1:1,000,000 | 1:10,000-1,000,000 | 1 |
| L3 | N42 | 0 | 0 | 1:100 |
| A3 | N42 | 0 | 1:10,000 | 1:100 |
| S3 | Bexa | 0 | 1 | 1 |

Isolates were grown on an overhead rotary shaker at 20°C for 48 hours in the presence of 8% or 2% of DMSO and the indicated amount of drug and plated in serial dilutions of 1:100. Estimates are by comparison with a negative control without drug.

“0” indicates no growth at all or colony diameters less than half of diameters of control at time zero, indicating severe growth defects. 1 indicates no apparent difference versus negative control. Numbers indicate inverse of number to multiply with to achieve density comparable to negative control. Note that in the case of isolate A1 Bexarotene, against the previous evidence collected, showed ineffective. The experiment was repeated using 10; 5 and 2.5 µM Bexarotene at 5% DMSO (Bexa-#3 and Bexa-#4). The difference in estimated efficacy versus control is due to a difference in incubation time between Bexa-#3 and Bexa-#4.

